# Attention and Retention Effects of Culturally Targeted Billboard Messages: An Eye-Tracking Study Using Immersive Virtual Reality

**DOI:** 10.1101/2024.09.10.610975

**Authors:** Moonsun Jeon, Sue Lim, Maria K. Lapinski, Stephen A. Spates, Gary Bente, Ralf Schmälzle

**Affiliations:** Department of Communication, Michigan State University, 404 Wilson Rd., East Lansing, 48824, MI, USA

**Author notes:** Corresponding Author. — Center for Avatar Research and Immersive Social Media Applications — Department of Communication, Michigan State University.

**Keywords:** cultural targeting, virtual reality, billboard paradigm, eye tracking, memory

## Abstract

Targeting, the creation of a match between message content and receiver characteristics, is a key strategy in communication message design. Cultural targeting, or adapting message characteristics to be congruent with a group’s cultural knowledge, appearance, or beliefs of recipients, is used in practice and is a potentially effective strategy to boost the relevance of a message, affecting attention to messages and enhancing effects. However, many open questions remain regarding the mechanisms and consequences of targeting. This is partly due to methodological challenges in experimentally manipulating messages that match cultural recipient characteristics while simultaneously measuring effects and balancing experimental control and realism. Here, we used a novel VR-based paradigm in which participants drove along a virtual highway flanked by billboards with varying message designs. Specifically, we manipulated the message design to either match or mismatch peoples’ cultures of origin. We used unobtrusive eye tracking to assess participants’ attention (i.e., for how long and how often they look at matched vs. unmatched billboards). Results show a tendency of the participants to inspect culturally matched billboards more often and for longer. We further found that matched billboards produce better recall, indicating more efficient encoding and storage of the messages. Our results underscore the effectiveness of cultural targeting and demonstrate how researchers can rigorously manipulate relevant message factors using virtual environments. We discuss the implications of these findings regarding theories of cultural targeting and methodological perspectives for the objective measurement of exposure factors through eye tracking.

## 1. Introduction

Imagine you are in a foreign country, all alone, and everything looks different from what you are used to. You notice a billboard that depicts a scene from your home country. You look at it more closely, keep inspecting it for a while, and remember it long after the end of your voyage. What are the reasons behind these effects? How do messages that match our cultural identity attract attention and promote retention? Cultural tailoring of messages is a method for matching information in persuasive messages to the cultural characteristics of potential receivers under the assumption that it will result in a greater message impact. Nonetheless, questions remain about the outcomes and mechanisms of such messages especially in terms of gains in attention and information retention.

In this paper, we first discuss the role of cultural targeting as a messaging strategy and review the ways it has been used in communication including explanations for its effects. Next, we will zoom in on the nexus between exposure and reception, or how messages available in peoples’ visual communication environments garner attention. We will point out that this exposure-reception nexus, which forms the theoretical foundation of message effects (Slater, 2004), has remained challenging to measure at the individual level, largely due to methodological bottlenecks. We argue that recent advances in virtual reality (VR) technologies with integrated eye tracking facilities create new opportunities to study the effects of cultural targeting with experimental rigor and precision and ecologically validity. We will then present the current study in which we display billboard messages presented in a realistic driving environment that either do or do not match participants’ racial identity and examine the effects of this manipulation on attention and subsequent retention.

### 1.1. Background

Messages congruent with receiver characteristics are likely to have greater resonance with receivers than those that do not (Kreuter & Wray, 2003; O’Keefe, 2002). Cultural targeting refers to the process of creating messages to address the cultural characteristics of receiver groups. This strategy is widely used in health communication, environmental awareness messaging, and persuasion at large (Cho, 2011; Huang & Shen, 2016; Joo & Liu, 2020). The basic idea behind cultural targeting is connected to the concepts of message targeting and tailoring broadly, which refer to messages designed for groups or individuals, respectively (Noar & Harrington, 2016). Cultural targeting differs from general message targeting in that it focuses on culturally-based differences regarding how groups might respond to messages (Lapinski & Oetzel, 2021). Culture, in this context, can be defined as a number of multi-faceted characteristics shared by a group (Kreuter et al., 2003).

Cultural targeting is believed to affect how people attend to,
receive, and process messages and ultimately their effects. Cultural targeting can take different forms and these forms are believed to change the impact of the messages on receivers (Kreuter & McClure, 2004; Huang & Shen, 2016); the forms range from those that target surface-level elements of culture to those that address deep cultural values (Resnicow et al., 1999). Lapinski and Oetzel (2021) expanded the work by Kreuter et al. (2003) to distinguish the core elements of message targeting and tailoring based on the literature on intercultural communication, health communication, and persuasion.

One surface-level element is that of design; these messages address surface-level elements of culture through the visual, verbal, and non-verbal content and involve “the matching of messages and intervention materials to the observable behavioral and social characteristics of cultural groups” (Lapinski & Oetzel, 2021, p. 250). Design elements can include messages which show people who look similar to potential receivers or involve exhibiting symbolic elements of culture (e.g., clothing, food, etc.). Researchers can also target more in-depth characteristics such as identity affiliation, values, or norms (Davis & Resnicow, 2012; Resnicow et al., 1999). More specifically, crafting messages that appeal to one’s social or cultural identity including ethnic, racial, gender, or other characteristics (e.g., Joseph et al., 2020) - is noted as one of the ways in which cultural targeting/tailoring can be implemented according to the cultural tailoring content (CTC) framework (Kreuter et al., 2003; Lapinski et al., 2024). In this study, we examine the effects of messages that match participants’ self-defined racial identity.

What can explain or predict the ways in which the elements of the design of messages shape message response? Theory development to explain the possible effects of cultural tailoring has been sparse, but there are several possible explanations for how and why design elements shape attention to messages and retention of the information in those messages. First, design elements of messages that show, for example, people who look like audience members may function to attract our attention based on principles of similarity such as mere exposure (Zajonc, 1968), liking or attraction (Simons et al., 1970), or the potential biological connections to people who look like us (Bailenson et al., 2008). That is, we tend to positively evaluate things that are familiar (Zajonc, 1968), like people who are more similar to us, and find them more socially attractive (Simons et al., 1970). We may even be biologically predisposed to think positively of people who are similar to us because it is evolutionarily functional (Bailenson et al., 2008). In each of these cases, our attention should be pulled toward objects and people that are more familiar or similar to ourselves relative to those that are less similar to us; when messages contain images of culturally similar others, we will be visually attracted to such images resulting in greater eye gaze and fixation.

Design elements that depict members of an ingroup with which one identifies might also operate at a deeper level. For example, racial identity matched messages might function to bring one’s social identity into focus. Social identity theory (SIT) explains how intergroup relations are perceived and unfold within interpersonal processes (Hornsey, 2008; Tajfel, 1974). SIT argues that people are inclined to establish social identity by making distinctions between their ingroup and outgroup, and by reinforcing them by attaching a positive image to their ingroup (Tajfel, 1978; Tajfel & Turner, 1979). Self-categorization theory (SCT) is related to SIT and postulates that individuals enhance their identity as a group when they are exposed to situations where their group membership becomes salient, whether it happens transiently or through a prolonged period of time and use it to inform their decision-making (Turner et al., 1987). Aligned with these theoretical assumptions, ingroup identification has been noted as a significant factor that affects message attention, recall, and subsequent persuasion effects, as exemplars of ingroup presented in messages are privileged to attract more attention compared to outgroup members (Hale, 2022; Hogg & Reid, 2006; Hogg & Smith, 2007; Wyer, 2010).

In short, there is reason to believe that design features of messages can harness basic psycho-social processes and as such can garner attention and improve retention. Empirical data suggest that messages that match a recipient’s cultural features, such as race, ethnicity, or values, are processed preferentially and have a larger effect (Huang & Shen, 2016) than untargeted messages, but it still warrants investigation on the mechanisms by which this advantage is achieved (Hawkins et al., 2008; Lapinski & Oetzel, 2021).

### 1.2. Examining Effects of Cultural Targeting: Old Challenges and New Opportunities

Although targeting, including cultural targeting, is one of the message strategies used in health communication research and practice, considerable questions remain regarding its underlying mechanisms in part because studies in public health, where cultural targeting is most often studied, typically tailor along multiple dimensions (Torres-Ruiz et al., 2018). Studying the effects of culturally targeted messages is complicated as it requires experimental designs in which messages are adapted to match specific individual or group characteristics, as well as measuring the minutiae of message reception mechanisms. Moreover, as with all communication and message effects research, it is a challenge to balance experimental control and ecological validity (e.g., Levine, 2018; Miller et al., 2019).

Here, we present a new solution to these challenges that leverages a recently described VR paradigm (Anonymous et al., 2023). This paradigm uses VR to create a message reception context, i.e., an artificial, but realistic and naturally navigable communication environment. In this case, participants experience a virtual drive along a realistic highway with billboard messages. Specifically, they enter a 3D-photogrammetry version of a real-world highway (highway 50 near Cold Springs, Nevada USA) that was digitized by the Nevada Department of Transportation, and they can drive through this environment much like in a driving simulator or an immersive computer game, but with the difference that they are actually fully immersed in the spatial surrounding.

One of the main benefits of using VR technology in message effects research is its realism (Martingano & Persky, 2021). Specifically, VR-immersed participants can drive along the highway at their own pace, look around, and explore their surroundings, which include various billboards that are typical of the US highway system. Unbeknownst to participants, who encounter these billboards as if they were just there, the messages that are placed on the billboard stands can be experimentally manipulated - a little bit like in the famous movies such as The Truman Show (1998), Minority Report (2002), or The Matrix (1999). This allows, for instance, to display messages that either match or don’t match participants’ racial identity (or any other message characteristic that is of theoretical interest to experimenters).

In addition to offering a way to create artificial but realistic communication environments that can be experimentally manipulated, another benefit of VR is the enormous measurement potential it affords. Specifically, via VR-integrated eye trackers, it becomes possible to keep track of where in their visual field participants are looking and relate this information to the information content in the environment, such as the specific billboard and whether it is targeted/matched or nontargeted/unmatched. Accurately measuring message exposure is crucial to all health intervention studies (Morris et al., 2009), and to communication in general (e.g., Slater, 2004). Thus, via a VR-integrated eye tracker, we can study how participants attend (i.e., look at) incidentally to messages as opposed to screen-based eye-tracking paradigms in which message exposure is typically forced. As such, VR is a methodological innovation for message effects research that promotes method-theory synergy because it allows researchers to manipulate features of messages and environments, rigorously measure relevant variables (as opposed to retrospective self-report or aggregate metrics of exposure), and do so in a way that resembles real-life visual information environments.

### 1.3. The Current Study and Hypotheses

The VR-based approach described above makes it possible to focus on the nexus between incidental exposure to messages (by letting participants drive and look freely), message reception (by recording the number of fixations and total gaze duration towards the billboards), and subsequent effects (by measuring which of the billboards they remember). As participants drive down the highway, they are exposed to culturally-targeted billboard messages, including those matched or not matched to participants’ self-identified race. These messages feature a variety of health-related topics that are typical of highway billboards, such as buckling-up, distracted driving, drunk driving, as well as smoking, alcohol, and other health, risk, and safety-related topics. Using VR, we can measure whether, for how long, and how often they look at billboards they pass, which rigorously quantifies their overt attention toward each message. Unbeknownst to participants, the VR-environment is pre-programmed to adapt to the participants’ cultural identity, i.e., the specific billboards that are displayed are manipulated to match/mismatch with the participant/group characteristic - which is the key rationale behind cultural targeting. Once participants arrive at the end of the highway, we test their incidental memory for the messages - first via an unannounced free recall test and then via a recognition memory test.

Drawing upon the theoretical framework outlined previously, we propose that the mechanisms underlying the effects of culturally targeted exemplars in this study can be explained from aforementioned research on social identification and source similarity (Hayashi et al., 2018; McQueen et al., 2011; Tharp-Taylor et al., 2012; Turner & Reynolds, 2012). Through exposure to messages that highlight one’s racial identity (i.e., presenting those that look similar), it is more likely that culturally targeted messages will activate one’s ingroup heuristics, which will, as a result, attract greater attention. This, in turn, would lead to greater retention abilities such as recall and recognition. Thus, based on the theory and rationale laid out above, we formulate the following hypotheses and research questions:

**H1**. Culturally targeted messages will command greater attention (i.e., **a)** longer gaze duration and **b)** more fixations) compared to non-targeted messages (i.e., mismatching messages and neutral messages).

**H2**. Culturally targeted messages will result in better retention (i.e., **a)** recognition and **b)** recall) compared to non-targeted messages (i.e., mismatching messages and neutral messages).

**H3**. Overt attention (i.e., **a)** greater fixations and **b)** longer gaze durations) will mediate the effect of culturally targeted messages on retention (i.e., recognition and recall), such that culturally targeted messages will lead to greater attention, which in turn leads to greater retention.

**RQ1**. How do different messages on billboards impact viewers’ attention and retention, and how do personal characteristics influence these outcomes?

## 2. Method

### 2.1. Participants

Forty participants (*M*_*age*_ = 22, *SD*_*age*_ = 6.2) were recruited from a university participant pool and via word of mouth. The eligibility criteria included 1) those over age 18 and 2) those who had a normal or adjusted vision. The sample size was determined a priori based on prior research (Anonymous et al., 2023). The study was approved by the university institutional review board, and all participants provided written informed consent and were compensated for their participation. We collected data from 40 participants and immediately replaced participants who reported blurry vision or experienced other issues (*n* = 2). We excluded one participant who identified as neither White Caucasian nor African American/Black. We also excluded one outlier after an exploratory analysis of the eyetracking metrics whose data was abnormal from the rest of the data. Therefore, the final sample size was 38^2^.

### 2.2. Materials and Equipment

#### Equipment and VR-based Eye-Tracking

The experiment was run in a VR research lab on a GPU-powered gaming PC. We used the HP Reverb G2 Omnicept VR headset with eye-tracking capabilities (Tobii AB, 120 Hz sampling rate). A python-based VR research platform (Vizard, Worldvision Inc.) was used to program the experiment, using the Sightlab VR package to capture eye-tracking in the VR environment.

The environment used for driving was a virtual 3D-photogrammetric model of a highway in Nevada that has also been used in prior research (Anonymous et al., 2023), in which VR-immersed participants could drive down while freely looking at billboard messages that were placed along the road (see Figure 1).

**Figure 1:**
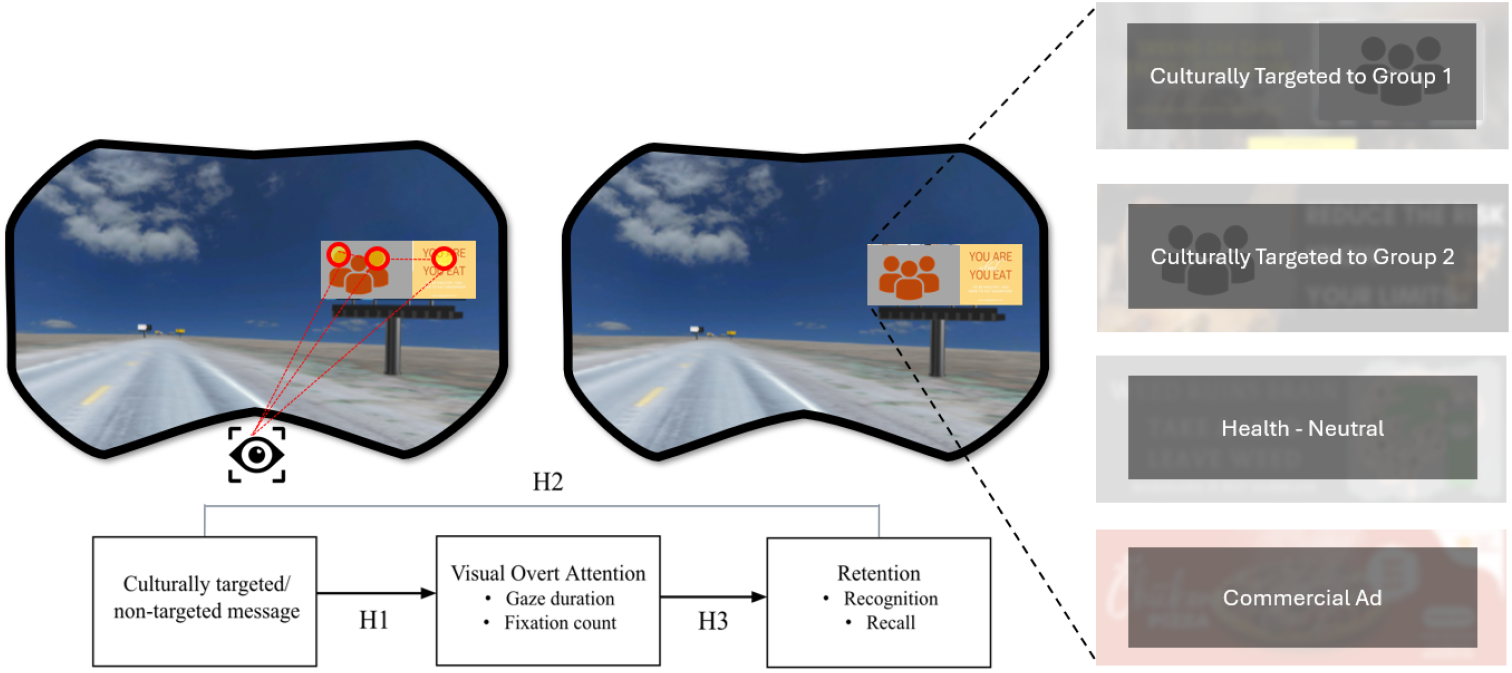
Illustration of the VR Billboard Paradigm and its adaptation in the current study to examine the effects of cultural targeting. A realistic photogrammetry-based model of a highway (Highway US50 in Nevada) serves as the virtual environment. Via a VR-integrated eye tracker, we can objectively track whether, for how long, and how often participants look at billboards they pass during a simulated drive. Unbeknownst to participants, the billboards are manipulated to either match or mismatch participants’ cultural identity. For instance, in the depicted exemplar, there are two versions depicting a family dinner among either a family of white/Caucasian Americans or Black/African Americans. In essence, each billboard exists in multiple versions (e.g. for buckling up, smoking prevention etc.) that attempt to maximize the overlap between participants’ cultural background and the way the message depicts models with a matching background. Our experimental hypotheses then focus on the attention (gaze behavior) towards the matched/unmatched billboards and the effect of matching on retention (memory). We obscure select examples of the original message stimuli to safeguard image rights.

#### Billboard Messages

Participants were exposed to a total of 20 billboard messages in their VR driving experience including 15 health messages and 5 commercial advertisements as fillers. 15 health messages covered a diverse range of topics, including binge drinking, underage drinking, having healthy eating habits, smoking, cannabis use, avoiding texting while driving, etc. Among the 15 health-related billboards the participants were exposed to, five billboards presented exemplars that matched their self-reported racial identity (i.e., culturally targeted), and another five presented exemplars that did not match their racial identity (i.e., culturally non-targeted). The placement, colors, and font of the texts and images were held constant throughout the two conditions; the only variation was the portrayed racial identity of the exemplars. The remaining five displayed health messages that contained non-human objects and texts without exemplars and included promotions for a law firm, burger restaurant, furniture sales, etc. The order of the billboards was randomized for all participants. The images of the billboards were adapted from image search using Google and only used when they were free of use in terms of copyrights. Other images were generated using the Midjourney.com generative AI service.

### 2.3. Experimental Procedure and Conditions

The procedure for this study comprised an arrival phase, a VR setup and a short demo phase, the main experimental highway drive, and a post-drive interview that included an unannounced memory test and survey questions. In brief, the participants came into the lab in person, signed the consent form, and completed a brief vision test before putting on the VR headset. Once they put on the VR headset, the researchers executed the VR eye-tracking calibration routine and then allowed the participants to practice navigating the demo virtual highway environment without the billboards. Next, the participants drove straight down the full virtual highway with the billboards. The full drive took about 10 minutes.

During the post-experiment phase, the participants were given a sudoku puzzle for 2 minutes to clear out their immediate short-term memory contents and prevent recency effects. Then, the researcher(s) conducted a brief structured interview, during which they asked the participants about the length of the drive, their general experience in VR, and every billboard-related factor they recalled (e.g., “Now, please tell us everything you remember about the billboards you passed.”). Next, they partook in the recognition test, marking the billboards that they recognized (shown on a paper survey along with multiple distractors). Finally, the participants completed an online survey via Qualtrics, which included questions about demographics and VR experience.

### 2.4. Measures

The main measured variables in this study comprise eyetracking data (whether/how long each billboard was looked at) and memory outcome measures (whether a billboard was later recalled or recognized; Kim & Southwell, 2017). Specifically, using the VR-integrated eye-tracker, we determined the total duration of gaze (i.e., for how long a given billboard was inspected by each driver). To assess memory, we used a free recall task administered at the end of the highway drive right after participants removed the VR headset and completed a short interview. Recognition memory was then tested via a pen and paper survey using a digital tablet that included all experimental billboard images along with distractor images. The same postexperimental survey also included measures of participants’ racial/cultural identity, demographic variables, as well as potential VR-related symptoms (Figure 2) and perceptions of spatial presence (adapted from Anonymous et al., 2023; Kim, 2023; Souza et al., 2021).

**Figure 2:**
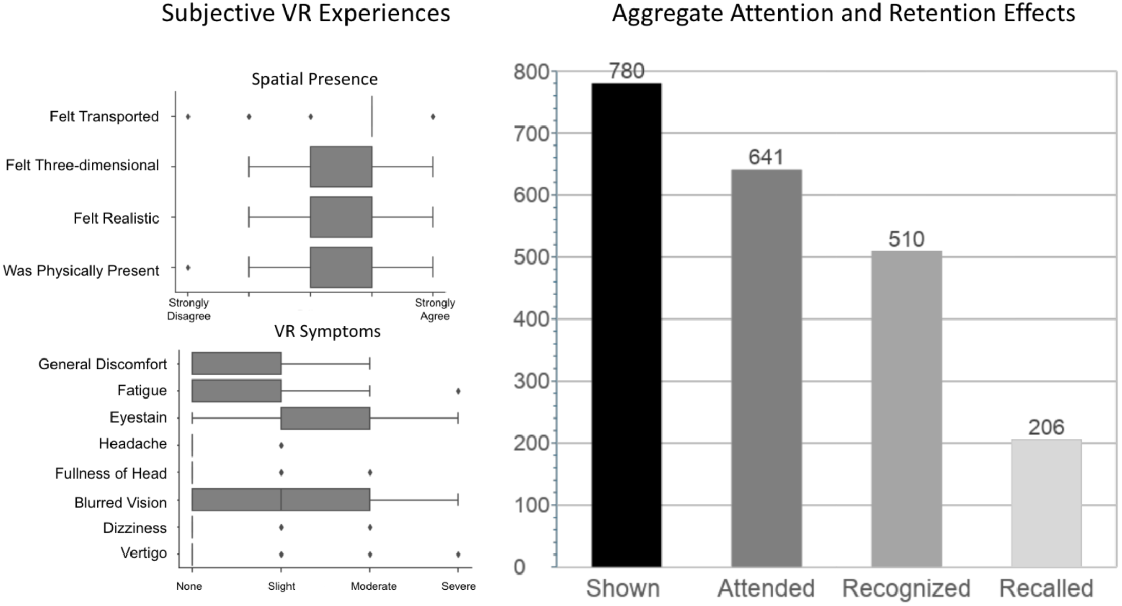
Left: Subjective reports about their experiences during the VR-mediated highway drive confirm that participants felt spatially present in the highway environment and experienced little to no symptoms. Right: Aggregate Effects of Exposure to Messages on Attention and Retention. Out of the 780 hypothetical exposures (38 participants all passing 20 billboards: 38*20 = 760), 641 billboards were explicitly looked at (fixated at least 1*), and 510 were recognized and 206 recalled in the post-drive interview. These data confirm the classical arguments of McGuire and advertising researchers more broadly who argue for a multiplicative decay-factor of ca. .7 along the exposure-to-attention-to-retention pathway.

### 2.5. Data Analysis

The main variables of our experiment were i) participants’ self-reported racial identity (person variable), ii) the content of the billboard (matching or non-matching or not relevant to racial identity), iii) how often participants looked at (fixated) on each billboard message and for how long they looked at it in total (self-defined behavioral variables), and iv) whether participants recalled/remembered the billboard (also self-defined). Importantly, while participant identity and billboard content were manipulated quasi-experimental or experimental variables, the viewing behavior and the message memory are self-defined behavioral variables (i.e., different participants decided to look at specific messages and not others, and different participants remembered different billboards).

Eye movement behavior and memory data were analyzed using Python code. We document the analysis and provide code in the study’s online repository at [anonymized for review]. Specifically, we read in the output file from the VR-eye-tracking, which contained information about which billboards a participant had fixated with the information from the post-study interview and online survey, where memory was assessed for all billboards. Thus, the analytical dataset comprises a table for each participant, containing information about which billboard and which version (e.g., a don’t text and drive billboard featuring African American models) was displayed, whether it was looked at, and whether it was subsequently remembered. Based on this dataset, we then collapsed data based on whether or not the message matched the participant’s racial identity and computed dependent variables in the resulting cells (e.g., share of fixated billboards in the matched vs. unmatched, share of recalled and remembered billboards in these conditions).

In the analysis, we first compared the effect of message-matching on attention (fixation count and gaze length) and retention (recall and recognition rate) separately and then combined both eye-tracking and memory outcomes in a joint analysis.

## 3. Results

### 3.1 Participants’ Experiences and Behavior During the VR Highway Drive

First, to examine participants’ experiences of spatial presence and the effects of VR during the drive, we analyzed their open-ended responses from the post-drive interview and survey answers. Overall, participants expressed that they felt immersed in the environment as if they were actually driving on the highway, and several participants described their driving experience as calm, and relaxing. Confirming these impressions, they reported above notably high levels of perceived spatial presence (*M* = 3.62, *SD* = 0.65; range 1–5) and below low levels of VR-related symptoms, such as dizziness or discomfort (*M* = 1.59, *SD* = 0.62; range 1–4) (Figure 2).

Next, we turn to participants’ behavior during the drive, examining whether they looked at the billboards. We found that many participants looked at almost all billboards they passed (*M* = 16.4, *SD* = 4; out of 20 possible), with only 10 participants looking at less than 15 of the 20 they passed, which could be due to participant inattention or measurement issues. In any case, these results confirm that participants scanned the dynamic visual environment they found themselves in and paid at least basic attention to their surroundings.

Across the 38 participants, each of whom passed 20 billboards (i.e., 38*20 = 760 “assumed” exposures), we obtained fixations on 641 billboards, or about 84% (“attended” exposure, cf. Southwell et al., 2002). Overall, participants recognized 510 billboards (67% of all possibilities and 80% of the attended ones), and they freely recalled 206, which amounts to about 27% (of the 760 presented) or 30% (of the 641 attended) billboards. Broadly, this funnel-shaped reduction from exposure to a minimal message effect - defined here as being able to actively retrieve the message from memory - is compatible with McGuire’s famous message effects framework (McGuire, 2011) and results from the advertising literature more broadly (Rossiter & Bellmann, 2005). Moreover, the survey results demonstrate that the participants experienced the drive as realistic, were not adversely affected by the VR technology, and reported a general sense of satisfaction and enjoyment (see Figure 2).

### 3.2. Effect of Cultural Targeting on Overt Visual Attention

#### Gaze Duration

Next, we examined the effect of matching/non-matching the billboard’s models to the participants’ cultural identity. To this end, we first examined whether matching affected the gaze duration of the billboards. Thus, for every participant, we computed one gaze duration measure for the matched billboards and a corresponding measure for the non-matched ones, and we compared these measures at the group level via a paired samples t-test. The results are illustrated in Figure 3 (top row). On average, participants looked at the messages that matched their cultural identity for about 3.074 seconds (*mean*_*GazeDuration: matched*_ = 3.074, *SD* = 1.493) while they gazed at the unmatched messages for only about 2.851 seconds (*mean*_*GazeDuration: unmatched*_ = 2.851, *SD* = 1.422). Although this suggests a nominally longer gaze duration towards matched messages, this difference was not statistically significant (*t*(37) = 1.213, *p* = .116). Furthermore, we assessed gaze duration towards neutral health messages (i.e., messages without human exemplars), which was, on average, about 3 seconds (*mean*_*GazeDuration: neutral health messages*_ = 3.135, *SD* = 1.246).

**Figure 3:**
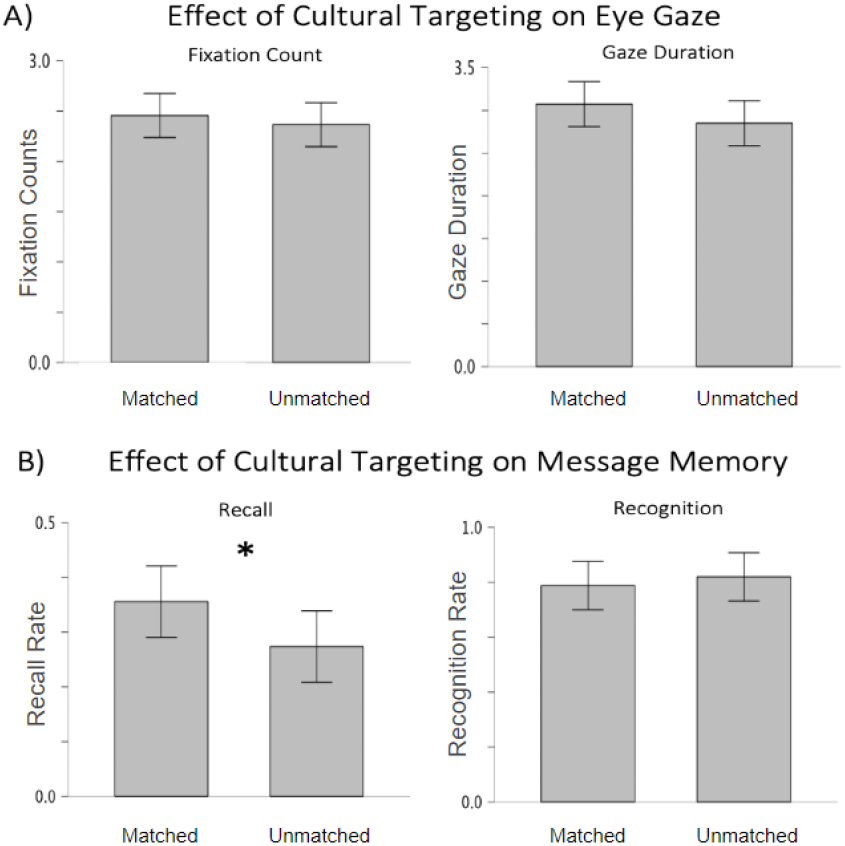
Top Row: Effect of Matching on Attention: Fixation Count and Gaze Duration. Bottom Row: Effect of Matching on Retention: Recall and Recognition

##### Fixation Count

The gaze duration measure above represents the total time spent inspecting a billboard. An alternative metric of visual attention to the billboards is the number of fixations towards a given billboard, meaning that a person might fixate on a billboard, then look somewhere else, but then revisit it for deeper re-inspection. As for the gaze duration measure, we found a slightly higher number of fixations towards matched compared to unmatched messages (*mean*_*FixationCount: matched messages*_ = 2.457, *SD* = 1.043; *mean*_*FixationCount: unmatched messages*_ = 2.365, *SD* = 0.959), but the statistical comparison was also not significant; *t*(37) = 0.601, *p* = .276). Again, we also assessed the number of fixations towards non-exemplar messages (i.e., neutral billboards about health-related topics without characters), finding them to lie at 2.542 fixations on average(*mean*_*FixationCount: neutral health messages*_ = 2.542, *SD*= 0.933).

Overall, although the pattern of results aligns with our prediction of more attention toward matched messages, the data are not consistent with H1.

### 3.3. Effect of Cultural Targeting on Retention

#### Recall

Having investigated the effect of cultural targeting on visual attention, we next investigated the effect of cultural targeting on retention. As expected, results showed that culturally targeted messages had a greater recall rate (*mean*_*Recall Rate: matched messages*_ = 0.356, *SD* = 0.249), compared to exposure to non-targeted messages (*mean*_*Recall Rate: unmatched messages*_ = 0.274, *SD* = 0.214). This difference was statistically significant (*t*(37) = 1.8, *p* = .040). For comparison, the recall rate for neutral health messages was again approximately similar to the unmatched messages, with a (*meanRecall* _*Rate: neutral health messages*_ = 0.297 (*SD* = 0.171). These results support H2a, which predicted better recall for targeted messages relative to untargeted messages.

#### Recognition

Running the same set of analyses for the data from the recognition task revealed the following results: As expected, the recognition rates are considerably higher than recall, with values of about 80% for recognition compared to 30% for free recall. Levels of recognition memory did not significantly differ between targeted and nontargeted messages (*mean*_*Recognition Rate: matched messages*_ = 0.788, *SD* = 0.262; *mean*_*Recognition Rate: unmatched messages*_ = 0.820, *SD* = 0.259; *t*(37) = -0.52, *p* = .697). As a comparison, the neutral health messages were recognized about 70% of the time (*mean*_*Recognition Rate: neutral health messages*_ = 0.788, *SD* = 0.262).

**Table 1.**
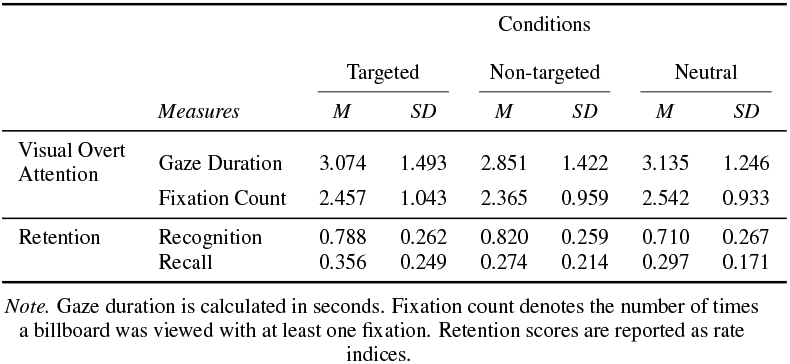
Means and Standard Deviations for Attention and Retention Measures Across Message Conditions.

| Measures |  | Conditions |  |  |  |  |  |
| --- | --- | --- | --- | --- | --- | --- | --- |
|  |  | Targeted |  | Non-targeted |  | Neutral |  |
|  |  | M | SD | M | SD | M | SD |
| Visual Overt Attention | Gaze Duration | 3.074 | 1.493 | 2.851 | 1.422 | 3.135 | 1.246 |
|  | Fixation Count | 2.457 | 1.043 | 2.365 | 0.959 | 2.542 | 0.933 |
| Retention | Recognition | 0.788 | 0.262 | 0.820 | 0.259 | 0.710 | 0.267 |
|  | Recall | 0.356 | 0.249 | 0.274 | 0.214 | 0.297 | 0.171 |

*Note.* Gaze duration is calculated in seconds. Fixation count denotes the number of times a billboard was viewed with at least one fixation. Retention scores are reported as rate indices.

### 3.4. Exploratory Analyses

In addition to testing our main a-priori hypotheses, we ran a series of exploratory analyses to address our research question. First, given the longstanding debates around single-vs. multiple-message designs (e.g., Clark, 1973; Jackson & Jacobs, 1983; O’Keefe, 2024), we zoomed in on gaze and memory results for individual messages. Second, we examined each racial subgroup of participants (i.e., Black/African American and White/Caucasian participants).

#### By-Message (By-Billboard) Analyses

Along with examining the average effects of matched and unmatched messages on attention and memory metrics, we examined the effects of matching on individual messages. To this end, we averaged the relevant metrics on a by-billboard basis and computed correlations between measures, as well as individual examinations of attention and retention effects. As can be seen in Figure 4, there were fairly evident differences across message conditions. In particular, among the 20 experimental messages, some were recalled (and also recognized) very well, whereas others achieved low recall and substantially lower recognition. When computing various vector-correlations between recall and recognition metrics, we find all those correlations to be positive (although not all were statistically significant because of the low number of items per cell, i.e., 10 targeted, of which 5 were matched/unmatched, 5 neutral/untargeted health messages, 5 untargeted commercial messages). Similar results were obtained for the attention metrics (fixation count and total gaze duration). These results suggest that there are inherent differences between the billboards, which affect interest and memorability. Given that we invested significant effort in controlling the billboards’ appearance and placement, we believe that these outcomes are due to topical factors (e.g., the alcohol messages being more relevant than, say furniture). Also, these results align with our prior findings (Anonymous et al., 2023), which also presented by-message differences. In fact, one common message across all studies - a billboard for a burger - was among the most highly recalled messages in this study. We speculate that the advertisement for a burger restaurant was too distinct from other sets of health message billboards.

**Figure 4:**
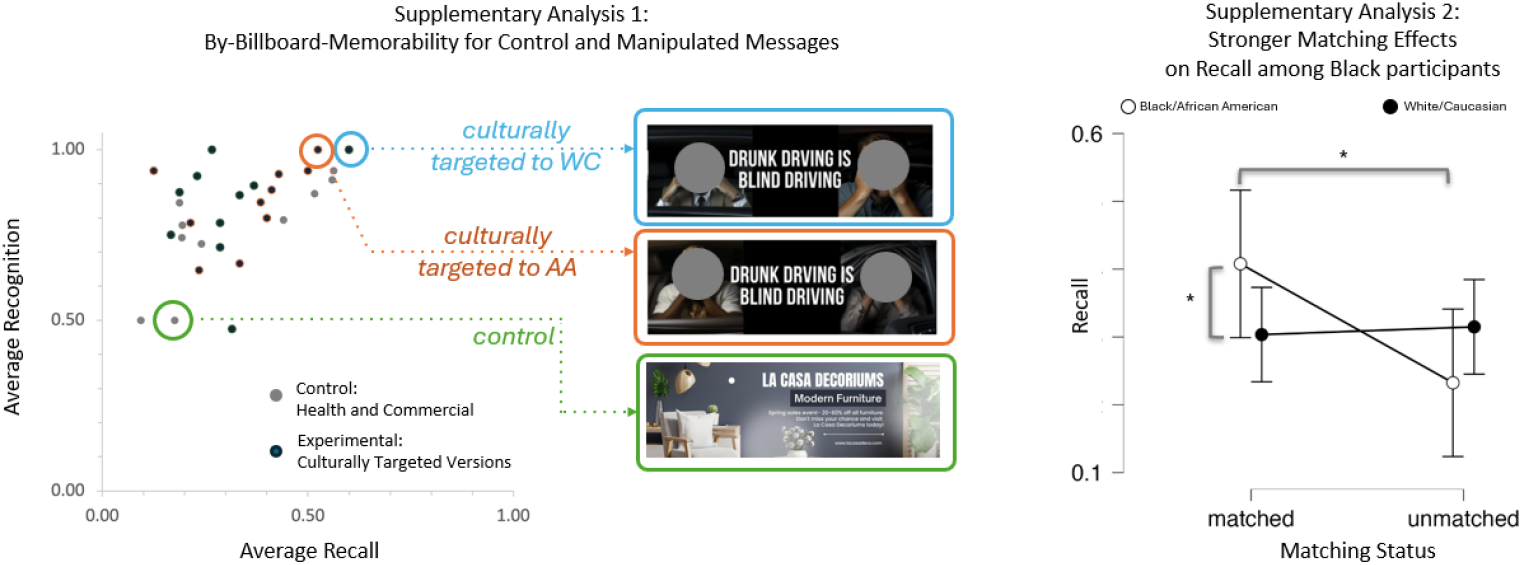
Illustration of Selected Results from Exploratory Analyses. Left panel: Examining By-Message (by-item/by-billboard) effects reveals robust correlations between individual billboards’ memorability, i.e., their average recall and recognition rates. Moreover, it is clearly evident that some billboards were rather memorable - and others not. The images used in the stimuli were generated by AI and do not raise any conflict with the copyright policy of the database. Right panel: In an exploratory analysis, we find an interaction between participants’ race and the cultural targeting effect in terms of message recall. Post-hoc analyses revealed that African American participants exhibit higher recall for culturally targeted (matched).

#### Analyses for Participant Subgroups (Culture/Race)

Examining the subgroups of participants revealed an interesting pattern of effects. First, for gaze metrics, we found significant main effects for self-identified race, which were driven by White/Caucasian participants generally looking longer at billboards and revisiting more billboards (For *Gaze Duration*: *F*(1,36) = 13.53, *p* <.001; For *Fixation Count*: *F*(1,36) = 6.5, *p* = .015). A parallel pattern was observed for recognition, with White/Caucasian participants having higher recognition rates compared to African American/Blacks (*Recognition*: *F*(1,36) = 5.7, *p* = .020). For recall, the results revealed an interesting interaction between matching status and participant race (*Recall*: *F*(1,36) = 4.6, *p* = .038). In particular, as illustrated in Figure 4 (right panel) and confirmed by post-hoc analysis, the African American/Blacks group showed a stronger matching effect, recalling more matched than unmatched messages (Recall matched >unmatched: *t* = 2.853, *p*_*holm*_ = .043).

## 4. Discussion

In this study, we examined the effect of cultural targeting of messages on people’s attention and memory via eye-tracking metrics within a virtual environment. We found that messages with images aligned with people’s self-reported racial identity were slightly more likely to draw overt visual attention (measured via fixation count and gaze duration), though this was not statistically significant across all messages. Furthermore, we found that people recalled culturally matched messages more than unmatched messages.

First, we discuss the effects of cultural targeting (matching) on visual attention. Our statistical analyses (using mixed-effects modeling to take into account variable message effects; Jackson & Jacobs, 1983) failed to identify strong effects of matching on visual attention. Specifically, we find that almost all of our participants fixated on each billboard at least once, suggesting a high base level of attention. Next, we zoomed in on the more nuanced attention metrics of gaze duration (how long they looked at the billboards in total) and fixation count (how often they revisited a given billboard). We found that both metrics do, in fact, show some evidence of more attention towards matched messages - gaze duration and fixation count were both nominally higher for the matched billboards compared to unmatched ones. However, this difference was not statistically significant. It remains speculative whether all of our message manipulations were explicit enough for participants to recognize the similarity of racial ingroup members portrayed on the billboards. In sum, these results do not support H1, although the by-message analyses (see Figure 4) suggested that the matching manipulation worked well for some messages.

Next, our second hypothesis stated that matched messages would lead to better retention. Our results provided support for this hypothesis, albeit with some qualification. We did find that matched messages were recalled significantly more often, which supports H2. However, for the recognition measure of memory, we did not find corresponding retention benefits for message matching. It should be noted that recognition memory in this context may not be the best measure of retention. Specifically, as shown in Figure 4, the messages all looked very similar, conflating the differences between the matched and unmatched messages. Our recognition test displayed only the versions of the messages to which participants were exposed and distractors to account for general guessing tendencies; it is possible that participants just “recognized” the message topic (e.g., buckling up), but not the specific billboard. In other words, they could have mixed up the different versions of the billboards, instead relying on an overall gist memory (e.g., Reyna & Brainerd, 1995; a billboard about buckling up with people on it) rather than the specific version of the billboard (e.g., two black people in a car accident). With this in mind, we are inclined not to place too much weight on the recognition measure and instead focus on message recall. In memory research, it is well-known that recall is a more difficult task because it requires the active retrieval of a memory trace, and we do find that matched messages were recalled more often. This is in line with our predictions and thus provides support for H2.

The observed advantage of matched messages in memory (free recall), yet the lack of prioritized attention (gaze behavior) towards them during encoding, leads to the question as to when/where in the processing stream this effect arises. If all messages were looked at (attended to overtly) and there were no big differences in either the duration or the frequency of looking at matched vs. unmatched messages (i.e., no support for H1), this suggests that the improved recall for matched messages must originate from a post-perceptual processing boost. Moreover, this effect is slightly at odds with previous work that found prioritized overt attention (in terms of eye gaze) was directly related to improved retention. We consider it likely that differences in the experimental setup explain these seemingly incompatible effects; specifically, in Anonymous et al. (2023), we instructed participants to perform a parallel, attention-consuming task and found that this task had a strong effect on overt attention (gaze behavior). However, in the current study, we only had a single “free viewing” task in which participants were free to explore the highway and its surroundings (including the billboards). As demonstrated by the fact that all participants looked at all billboards, this has led to a ceiling effect in terms of attention deployment towards the visual environment. Thus, given that we can empirically verify that all participants had viewed all billboards, we concluded that the recall-advantage for matched billboards must come into play after the stage of overt attention.

The observed effect of culturally-matched messages on participants’ memory may be attributed to enhanced self-relevance, as suggested by Schmitz & Johnson (2007). Given that the billboards in the present study predominantly focused on diverse health issues, it might be the case that racial-identity matched exemplars heightened risk perceptions associated with the diseases depicted (Goldstein et al., 2021). For instance, when a person who self-identifies as Black is shown a billboard featuring a Black person suffering from neck cancer due to smoking, it may intensify their perceived susceptibility to the disease by emphasizing similarities between the viewer and the exemplar. When these culturally targeted messages are presented alongside culturally non-targeted messages, they might be more effective in promoting such risk perceptions as they are recalled better. This is also related to the own-race effect (ORE) in memory for faces, which demonstrates that people are inclined to remember faces of people who are perceived as their own race, compared to less familiar, or of other race faces (Brigham & Malpass, 1985; Meissner& Brigham, 2001; Zhou et al., 2021). However, in order for such an effect to be observed, it is important that detection between in-group and out-group is established in the visual field (Prunty et al., 2023). Since almost all participants fixated on every billboard in our study, it was difficult to parcel out the impact of such detection in the current study. Previous research on own-race face detection relies on reaction-time measures of accuracy and speed recognition (e.g., Barkowitz & Brigham, 1982; Wiese et al., 2014), whereas we used eye-tracking. Moreover, when it comes to understanding the mechanisms of own-race effect, other factors like attitudes and interracial contact might come into play (Meissner & Brigham, 2001) that go beyond the current data. Thus, while work on the own-race effect is relevant to the current study and research context, we believe more investigation is needed to integrate prior research with the current paradigm and cultural targeting framework.

Our supplementary analyses revealed additional insights worth mention. For instance, we found that different billboards varied in terms of the degree to which they were looked at and how often they were recalled. Methodologically, this underscores the importance of modeling message-level variability in statistical analyses (e.g., Jackson & Jacobs, 1983), but it also raises several questions, such as whether our experimental matching manipulation was equally effective for all billboard exemplars as well as whether different health topics are more or less easily recalled. Generally, the between-message differences also align with prior work in which we found similar differences between more easy-to-recall topics (like buckling up) and more obscure topics.

### 4.1. Broader Implications: Theoretical and Applied

This research constitutes a new effort to investigate the mechanisms of cultural targeting, gauging message recipients’ overt visual attention and retention in a virtual environment setting with high ecological validity compared to traditional laboratory experiments. Our findings corroborate the notion that targeting one’s cultural identity with specific message content impacts message recognition and recall (Hale, 2022; Nelson & Garst, 2005). Moreover, we were able to observe that these effects may be intensified among specific groups, such as African Americans/Blacks, which aligns with the previous empirical findings on cultural targeting/tailoring messages that presented similar results (e.g., Campbell & Quintiliani, 2006).

A recent review of culturally targeted/tailored messages notes a lack of theories that account for how and when culturally targeted/tailored message interventions should be most persuasive (Lapinski et al., 2024). According to Lapinski et al. (2024)’s review, only half of the cultural targeting studies actually examined social/cultural identity as the key targeting/tailoring component. In our study, the messages were certainly targeted, but questions remain whether this manipulation led to increased identification as opposed to merely familiarity-based effects for the targeted messages. In any case, this all raises the point that studies often use diverse definitions and operationalizations of cultural targeting/tailoring, and employ suboptimal measurement methods to demonstrate that these manipulations are effective, i.e., lead to more attention, stronger identification, or whatever the theorized mechanism may be. Although our study cannot resolve these issues completely, the use of objective measures of eye-tracking to investigate attention as a first and necessary precondition for stronger targeting/tailoring effects points the way to improved ways of studying cultural targeting’s mechanisms.

Moreover, this study adds an innovative methodological effort to the current cultural targeting/tailoring literature by operationalizing attention as part of the underlying process to explain message effects and gauging eye-tracking measures to empirically test such claims. While several existing research targets one’s cultural identity to test the message effects of culturally targeted messages, most rely on self-report survey methods, which might be vulnerable to response bias or memory distortions. By contrast, the present study uses eye-tracking method, which allows an unobtrusive measurement to objectively track overt attention as a key element of the message reception process, which in turn helps explain message effects (King et al., 2019). Additionally, the virtual reality billboard paradigm along with its within-subject design, provides high ecological validity. As our results indicate, participants are more likely to be immersed in the research environment that is created by the researchers, which enhances perceived realism compared to traditional laboratory studies (Fox et al., 2019), yet maintains the strengths of lab-based research such as high controllability and effective message manipulations.

Regarding the practical implications, this research offers a bright future for message designers, marketers, and public health communicators who are interested in testing the effectiveness of culturally targeted/tailored messages by applying emerging digital platforms and immersive technologies in diverse contexts. For instance, message communicators could collect more evidence to create engaging and memorable virtual billboards that resonate with specific cultural groups. Public health practitioners could utilize VR environments to test and refine culturally targeted/tailored messages before implementing large-scale interventions, and potentially advance message modalities by applying them in different digital platforms such as Metaverse. As communication interventions utilizing novel technologies and algorithms are becoming increasingly prevalent and advanced, even to the level of intricate personalization and tailoring, it is our hope that more communication scholars can attend to generating culturally-resonating messages that potentially aid various types of populations in different contexts.

### 4.2. Limitations and Strengths

Although the current study demonstrates the effects of cultural targeting and uses a promising VR-based paradigm, some limitations should be acknowledged. First, while VR generally offers the possibility to create infinitely flexible but realistic and well-controlled environments, the current choice of a highway driving context is only one singular instance of possible environments and may not maximize the potential of cultural targeting. Rather, it may be that targeting effects are more likely to be found in visually overloaded environments such as cities or other environments where the space is filled with “outgroup” information, which could make culturally matched billboards stand out more conspicuously (e.g., Wolfe & Horowitz, 2017). There could have been individual variability in terms of whether one uses race extensively as a metrics for ingroup and outgroup perceptions, as the impact of ingroup exemplars on social identification and categorization might be weighted differently among individuals (Fazio & Dunton, 1997).

Another limitation of this study relates to the strength of the experimental manipulation. In particular, Figure 4 shows that while the billboards are very clearly matched to specific target groups, different versions of each feature a significant degree of overlap. Thus, the targeting manipulation is a fairly small subset of the entire information contained in the billboard. For example, billboard features we could also have manipulated include the text or other characteristics (like color scheme or cultural symbols). However, manipulating these characteristics via targeting would have added many visual confounds. Therefore, we opted for a tighter experimental control. Despite this limitation, the fact that we were still able to see targeting effects (nominally on the gaze metrics and significantly for the recall) speaks to the potential persuasive power of targeting. Specifically, we argue that in the real world, where targeting manipulations are likely more subtle but also more powerful and particularly more spread out over time, the effects might accumulate to have a substantial impact. For example, in the current study, the recall benefit of matched messages was about 35% recall for matched messages compared to 30% for unmatched ones. On the face of it, this may seem like a relatively small advantage. However, if we consider that an average highway billboard might be passed by about 10000 cars per day, this would translate into 500 people more who would actively recall the message. Given that active recall is a reasonably high standard for the impact of messages, such effects matter greatly in real-world application (e.g., Percy & Rossiter, 1992).

### 4.3. Avenues for Future Research

This study provides several important directions for additional research. Especially as the metaverse develops, we might expect to encounter environments that can be preconfigured to maximize the overlap of environmental features with participants’ cultural identities, preferences, and other proclivities. In fact, it is common to show participants specific feeds and other ‘matching’ messages on social media platforms such as Facebook or YouTube. Though this is not exactly tailoring/targeting, it is easy to see the parallels and opportunities (or threats). Going forward, it might be possible to match the environment (e.g., the room, type of highway) and the message content (i.e., the models shown.) to participants’ cultural and interest-based characteristics.

Theory-building regarding the ways in which cultural targeting impacts message response can only proceed with additional research on ways in which the various message elements impact message response through McGuire’s hierarchy of effects. Indeed, crossing the table of message elements described recently in Lapinski et al. (2024) with the hierarchy of effects could result in a large number of testable hypotheses about the many possible message modifications and their impact. Message elements could be targeted (as we have done in this study) or tailored to individual cultural characteristics. Because the message elements range from surface to deep-level cultural targeting, examining the effects of the range of elements will be useful for identifying the mechanisms of message effects with a goal of prediction.

Methodologically, we have only scratched the surface of the information the eye-tracking (and behavior tracking more broadly) offers in these new communication environments. In the current study, we used two eye-tracking metrics gaze duration and fixation count - that represent fairly straightforward global metrics. For future studies, we could add the possibility of having specific areas-of-interest (AOIs) on the billboards (e.g., separate text and image information), or use various data-driven metrics and predictive modeling approaches to isolate the effective informational ingredients (e.g., human faces, fear/threat imagery, etc.) that drive experimental effect.

## 5. Summary and Conclusions

To summarize, we examined the effects of cultural targeting on participants’ visual attention and memory using the VR-based billboard paradigm. It was revealed that matched messages are preferentially recalled. As we enter the era of metaverse-mediated communication, it will become easy and likely commonplace to tailor content to recipient characteristics. From a communication intervention perspective, this raises new opportunities to examine the effects of messages that use cultural targeting in many applied areas. We can expect that targeting will continue to be used as a powerful strategy to influence people within risk, environment, and health communication-related areas.

## Acknowledgement

We thank Dana Anafina for her help during data collection.

## Data Availability

This study’s data are available on Github, at https://github.com/nomcomm/vr_billboard_t.

## Footnotes

2 The ratio of the self-identified racial identity of the participants was 50:50 (i.e., White (*n* = 19) and Black/African American (*n* = 19). Three participants from the African American/Black group identified themselves as mixed-race.

## References

Bailenson, J. N., Iyengar, S., Yee, N., & Collins, N. A. (2008). Facial similarity between voters and candidates causes influence. Public Opinion Quarterly, 72(5), 935–961. 10.1093/poq/nfn064

Barkowitz, P., & Brigham, J. C. (1982). Recognition of faces: Own-race bias, incentive, and time delay Journal of Applied Social Psychology, 12(4), 255–268. 10.1111/j.1559-1816.1982.tb00863.x

Brigham, J. C., & Malpass, R. S. (1985). The role of experience and contact in the recognition of faces of own- and other-race persons. Journal of Social Issues, 41(3), 139–155. 10.1111/j.1540-4560.1985.tb01133.x

Campbell, M. K., & Quintiliani, L. M. (2006). Tailored interventions in public health: Where does tailoring fit in interventions to reduce health disparities? American Behavioral Scientist, 49(6), 775–793. 10.1177/0002764205283807

Cho, H. (Ed.). (2011). Health communication message design: Theory and practice. Sage Publications.

Clark, H. H. (1973). The language-as-fixed-effect fallacy: A critique of language statistics in psychological research. Journal of Verbal Learning and Verbal Behavior, 12(4), 335–359. 10.1016/S0022-5371(73)80014-3

Davis, R. E., & Resnicow, K. (2012). The cultural variance framework for tailoring health messages. In H. Cho (Ed.), Health communication message design: Theory and practice (pp. 115–135). Sage Publications.

Fazio, R. H., & Dunton, B. C. (1997). Categorization by race: The impact of automatic and controlled components of racial prejudice. Journal of Experimental Social Psychology, 33(5), 451–470. 10.1006/jesp.1997.1330

Fox, J., Arena, D., & Bailenson, J. N. (2009). Virtual reality: A survival guide for the social scientist. Journal of Media Psychology, 21(3), 95–113. 10.1027/1864-1105.21.3.95

Goldstein, A. O., Jarman, K. L., Kowitt, S. D., Queen, T. L., Kim, K. S., Shook-Sa, B. E., Sheeran, P., Noar, S. M., & Ranney, L. M. (2021). Effect of cigarette constituent messages with engagement text on intention to quit smoking among adults who smoke cigarettes: A randomized clinical trial. JAMA Network Open, 4(2), e210045. 10.1001/jamanetworkopen.2021.0045

Hale, B. J. (2022). Examining the effect of identification with a social media community on persuasive message processing and attitude change. New Media & Society, 26(8), 4589–4610. 10.1177/14614448221124085

Hayashi, H., Tan, A., Kawachi, I., Minsky, S., & Viswanath, K. (2018). Does segmentation really work? Effectiveness of matched graphic health warnings on cigarette packaging by race, gender and chronic disease conditions on cognitive outcomes among vulnerable populations. Journal of Health Communication, 23(6), 523–533. 10.1080/10810730.2018.1474299

Hawkins, R. P., Kreuter, M., Resnicow, K., Fishbein, M., & Dijkstra, A. (2008). Understanding tailoring in communicating about health. Health Education Research, 23(3), 454–466. 10.1093/her/cyn004

Hogg, M. A., & Reid, S. A. (2006). Social identity, self-categorization, and the communication of group norms. Communication Theory, 16(1), 7–30. 10.1111/j.1468-2885.2006.00003.x

Hogg, M. A., & Smith, J. R. (2007). Attitudes in social context: A social identity perspective. European Review of Social Psychology, 18(1), 89–131. 10.1080/10463280701592070

Hornsey, M. J. (2008). Social identity theory and self-categorization theory: A historical review. Social and Personality Psychology Compass, 2(1), 204–222. 10.1111/j.1751-9004.2007.00066.x

Huang, Y., & Shen, F. (2016). Effects of cultural tailoring on persuasion in cancer communication: A metaanalysis. Journal of Communication, 66(4), 694–715. 10.1111/jcom.12243

Jackson, S., & Jacobs, S. (1983). Generalizing about messages: Suggestions for design and analysis of experiments. Human Communication Research, 9(2), 169–191. 10.1111/j.1468-2958.1983.tb00691.x

Joo, J. Y., & Liu, M. F. (2021). Culturally tailored interventions for ethnic minorities: A scoping review. Nursing Open, 8(5), 2078–2090. 10.1002/nop2.733

Joseph, R. P., Keller, C., Vega-López, S., Adams, M. A., English, R., Hollingshead, K., Hooker, S. P., Todd, M., Gaesser, G. A., & Ainsworth, B. E. (2020). A culturally relevant smartphone-delivered physical activity intervention for African American women: Development and initial usability tests of smart walk. JMIR mHealth and uHealth, 8(3), e15346. 10.2196/15346

Kim, S. J. (2023). Virtual fashion experiences in virtual reality fashion show spaces. Frontiers in Psychology, 14, 1–13. 10.3389/fpsyg.2023.1276856

Kim, S., & Southwell, B. G. (2017). Memory for media content in health communication. In Oxford research encyclopedia of communication. 10.1093/acrefore/9780190228613.013.282

King, A. J., Bol, N., Cummins, R. G., & John, K. K. (2019). Improving visual behavior research in communication science: An overview, review, and reporting recommendations for using eye-tracking methods. Communication Methods and Measures, 13(3), 149–177. 10.1080/19312458.2018.1558194

Kreuter, M. W., Lukwago, S. N., Bucholtz, D. C., Clark, E. M., & Sanders-Thompson, V. (2003). Achieving cultural appropriateness in health promotion programs: Targeted and tailored approaches. Health Education & Behavior, 30(2), 133–146. 10.1177/1090198102251021

Kreuter, M. W., & McClure, S. M. (2004). The role of culture in health communication. Annual Review of Public Health, 25, 439–455. 10.1146/annurev.publhealth.25.101802.123000

Kreuter, M. W., & Wray, R. J. (2003). Tailored and targeted health communication: Strategies for enhancing information relevance. American Journal of Health Behavior, 27(3), S227–S232. 10.5993/AJHB.27.1.s3.6

Lapinski, M. K., & Oetzel, J. G. (2021). Cultural tailoring of environmental communication interventions. In B. Takahashi, J. Metag, J. Thaker, & S. E. Comfort (Eds.), The handbook of international trends in environmental communication (pp. 248–267). Routledge. 10.4324/9780367275204

Lapinski, M. K., Oetzel, J. G., Park, S., & Williamson, A. J. (2024). Cultural tailoring and targeting of messages: A systematic literature review. Health Communication, 1-14. 10.1080/10410236.2024.2369340

Levine, T. R. (2018). Ecological validity and deception detection research design. Communication Methods and Measures, 12(1), 45–54. 10.1080/19312458.2017.1411471

Martingano, A. J., & Persky, S. (2021). Virtual reality expands the toolkit for conducting health psychology research. Social and Personality Psychology Compass, 15(7), e12606. 10.1111/spc3.12606

McGuire, W. J. (1989). Theoretical foundations of campaigns. In R. Rice & C. Atkin (Eds.), Public communication campaigns (pp. 133–146). Sage.

McQueen, A., Kreuter, M. W., Kalesan, B., & Alcaraz, K. I. (2011). Understanding narrative effects: The impact of breast cancer survivor stories on message processing, attitudes, and beliefs among African American women. Health Psychology, 30(6), 674–682. 10.1037/a0025395

Meissner, C. A., & Brigham, J. C. (2001). Thirty years of investigating the own-race bias in memory for faces: A meta-analytic review. Psychology, Public Policy, and Law, 7(1), 3–35. 10.1037/1076-8971.7.1.3

Morris, D. S., Rooney, M. P., Wray, R. J., & Kreuter, M. W. (2009). Measuring exposure to health messages in community-based intervention studies: A systematic review of current practices. Health Education & Behavior, 36(6), 979–998. 10.1177/1090198108330001

Miller, L. C., Shaikh, S. J., Jeong, D. C., Wang, L., Gillig, T. K., Godoy, C. G., Read, S. J., & Akçayir, M. (2019). Causal inference in generalizable environments: Systematic representative design. Psychological Inquiry, 30(4), 173–202. 10.1080/1047840X.2019.1693866

Nelson, T. E., & Garst, J. (2005). Values-based political messages and persuasion: Relationships among speaker, recipient, and evoked values. Political Psychology, 26(4), 489–516. 10.1111/j.1467-9221.2005.00428.x

Niccol, A. (Writer, Producer), & Weir, P. (Director, Producer). (1998). The Truman Show [Film]. Scott Rudin Productions.

Noar, S. M., & Harrington, N. G. (2016). Tailored communications for health-related decision-making and behavior change. In M. A. Diefenbach, S. Miller-Halegoua, & D. J. Bowen (Eds.), Handbook of health decision science (pp. 251–263). Springer. 10.1007/978-1-4939-3486-718

O’Keefe, D. J. (2002). Persuasion: Theory and research (2nd ed.). Sage Publications.

O’Keefe, D. J. (2024). Message-level claims require message-level data analyses: Aligning claims and evidence in communication research. Communication Methods and Measures, 18(1), 1–14. 10.1080/19312458.2024.2349987

Percy, L., & Rossiter, J. R. (1992). Advertising stimulus effects: A review. Journal of Current Issues & Research in Advertising, 14(1), 75–90. 10.1080/10641734.1992.10504982

Prunty, J. E., Jenkins, R., Qarooni, R., & Bindemann, M. (2023). In-group and outgroup differences in face detection. British Journal of Psychology, 114(S1), 94–111. 10.1111/bjop.12588

Resnicow, K., Braithwaite, R., Ahluwalia, J., & Baranowski, T. (1999). Cultural sensitivity in public health: Defined and demystified. Ethnicity and Disease, 9(1), 10–21.

Reyna, V. F., & Brainerd, C. J. (1995). Fuzzy-trace theory: An interim synthesis. Learning and Individual Differences, 7(1), 1–75. 10.1016/1041-6080(95)90031-4

Rossiter, J. R., & Bellman, S. (2005). Marketing communications: Theory and applications (1st ed.). Prentice Hall.

Schmitz, T. W., & Johnson, S. C. (2007). Relevance to self: A brief review and framework of neural systems underlying appraisal. Neuroscience & Biobehavioral Reviews, 31(4), 585–596. 10.1016/j.neubiorev.2006.12.003

Simons, H. W., Berkowitz, N. N., & Moyer, R. J. (1970). Similarity, credibility, and attitude change. Psychological Bulletin, 73(1), 1–16. 10.1037/h0028429

Slater, M. D. (2004). Operationalizing and analyzing exposure: The foundation of media effects research. Journalism & Mass Communication Quarterly, 81(1), 168–183. 10.1177/107769900408100112

Southwell, B. G., Barmada, C. H., Hornik, R. C., & Maklan, D. M. (2002). Can we measure encoded exposure? Validation evidence from a national campaign. Journal of Health Communication, 7(5), 445–453. 10.1080/10810730290001800

Souza, V., Maciel, A., Nedel, L., & Kopper, R. (2021). Measuring presence in virtual environments: A survey. ACM Computing Surveys, 54(8), 1–37. 10.1145/3466817

Spielberg, S. (Director). (2002). Minority Report [Film]. 20th Century Fox.

Tajfel, H. (1974). Social identity and intergroup behaviour. Social Science Information, 13(2), 65–93. 10.1177/053901847401300204

Tajfel, H. (1978). Differentiation between social groups: Studies in the social psychology of intergroup relations. Academic Press.

Tajfel, H., & Turner, J. C. (1979). An integrative theory of intergroup conflict. In W. G. Austin & S. Worchel (Eds.), The social psychology of intergroup relations (pp. 33–47). Brooks/Cole.

Tharp-Taylor, S., Fryer, C. S., & Shadel, W. G. (2012). Targeting anti-smoking messages: Does audience race matter? Addictive Behaviors, 37(7), 844–847.

Torres-Ruiz, M., Robinson-Ector, K., Attinson, D., Trotter, J., Anise, A., & Clauser, S. (2018). A portfolio analysis of culturally tailored trials to address health and healthcare disparities. International Journal of Environmental Research and Public Health, 15(9), 1859–1872. 10.3390/ijerph15091859

Turner, J. C., Hogg, M. A., Oakes, P. J., Reicher, S. D., & Wetherell, M. S. (1987). Rediscovering the social group: A self-categorization theory. Blackwell.

Turner, J. C., & Reynolds, K. J. (2012). Self-categorization theory. In P. A. M. Van Lange, A. W. Kruglanski, & E. T. Higgins (Eds.), Handbook of theories of social psychology (pp. 399–417). Sage Publications Ltd. 10.4135/9781446249222.n46

Wachowski, L. (Director, Writer), Wachowski, L. (Director, Writer), & Silver, J. (Producer). (1999). The Matrix [Film]. Warner Bros.

Wiese, H., Kaufmann, J. M., & Schweinberger, S. R. (2014). The neural signature of the own-race bias: Evidence from event-related potentials. Cerebral Cortex, 24(3), 826–835. 10.1093/cercor/bhs369

Wolfe, J. M., & Horowitz, T. S. (2017). Five factors that guide attention in visual search. Nature Human Behaviour, 1(3), 1–8. 10.1038/s41562-017-0058

Wyer, N. A. (2010). Selective self-categorization: Meaningful categorization and the in-group persuasion effect. Journal of Social Psychology, 150(5), 454–470. 10.1080/00224540903365521

Zajonc, R. B. (1968). Attitudinal effects of mere exposure. Journal of Personality and Social Psychology, 9(2, Pt.2), 1–27. 10.1037/h0025848

Zhou, X., Burton, A. M., & Jenkins, R. (2021). Two factors in face recognition: Whether you know the person’s face and whether you share the person’s race. Perception, 50(6), 524–539. 10.1177/03010066211014016

